# Twist1 and balanced retinoic acid signaling act to suppress cortical folding in mice

**DOI:** 10.1101/2022.09.27.509818

**Authors:** Khue-Tu Ho-Nguyen, Manav Jain, Matt J. Matrongolo, Phillip S. Ang, Samantha Schaper, Max A. Tischfield

## Abstract

Evolution of cortical folding in gyrencephalic animals enabled higher cognitive functions and complex behaviors. Gene expression patterns and signaling molecules that control cortical folding have only recently been described and thus are still not well understood. In transgenic mouse models with induced cortical folding, amplification of neuroprogenitor cells or loss of their adhesion from the apical ventricular surface leads to gyri formation, whereas decreased cell adhesion in migrating projection neurons causes abnormal neuronal clustering and development of cortical fissures that resemble sulci. We now report that loss of *Twist1* expression in the primitive meninx results in cortical folding and sulci formation in the dorsolateral telencephalon. In developing sulcal regions, generation of apical and basal neuroprogenitor cells is normal. Instead, cell proliferation in the developing meninges is reduced, leading to loss of arachnoid fibroblasts that express Raldh2, an enzyme required for retinoic acid synthesis. Maternal retinoic acid supplementation rescues cortical folding and sulci formation. Our results suggest that balanced retinoic acid signaling from the meninges is required to maintain lissencephaly in mice, and in a manner independent from neuroprogenitor cell amplification.

## Introduction

Cortical folding occurs in a stepwise process centered upon the amplification of neuroprogenitor cell populations and heightened levels of neurogenesis (Llinares-Benadero and Borrell, 2019; Del-Valle-Anton and Borrell, 2022). Studies in mice, ferrets, and non-human primates show that increasing the number of apical neuroprogenitors (i.e., apical radial glia cells, aRGCs) in the ventricular zone (VZ) leads to amplification of basal radial glia (bRGC) and intermediate progenitor cells (IPs) (Chizhikov et al., 2019; Heide et al., 2020; Ju et al., 2016; Matsumoto et al., 2017; Rash et al., 2013; Roy et al., 2019; Stahl et al., 2013). Basal neuroprogenitors also have the ability to self-renew, which ultimately increases the numbers of mature projection neurons, causing folding. Gyrencephaly is generally assumed to have evolved from smooth brains (lissencephaly) (Welker, 1990). However, it is an evolutionarily labile trait as evidence suggests lissencephaly in mammals such as mice and marmosets arose from secondary loss of folding present in gyrencephalic ancestors (secondary lissencephaly), necessitated by decreases in body mass and other physiological adaptations (Kelava et. al., 2013). This suggests signaling mechanisms are required to actively maintain secondary lissencephaly in mammals.

Factors that promote secondary lissencephaly in mice have been recently identified and include non-cell autonomous BMP4 signaling from embryonic cranial mesenchyme and cell autonomous expression of adhesion proteins FLRT1 and FLRT3 in migrating projection neurons (Chizhikov et al., 2019; Del Toro et al., 2017). Loss of BMP4 signaling in *Lmx1a^−/−^/1b*^−/−^ embryos causes the formation of midline gyri via transient upregulation of Wnt/β-catenin signaling in the cortical hem (Chizhikov et al., 2019). This leads to an accumulation of aRGCs and subsequently, the generation of bRGCs and greater numbers of TBR2+ IPs. Conversely, *Flrt1^−/−^/Flrt3*^−/−^ animals develop cortical folding and sulci in the dorsolateral telencephalon through a process independent of progenitor cell amplification. Instead, loss of these adhesion proteins alters the radial migration of newborn projection neurons and causes abnormal neuronal clustering along the tangential axis of the cortical plate (Del Toro et al., 2017). Findings in *Lmx1a^−/−^/1b*^−/−^ and *Flrt1^−/−^/Flrt3*^−/−^ mutants, and those in *Trnp1^−/−^* mice which develop gyri via amplification of bRGCs (Stahl et al., 2013), suggest signaling processes that lead to expansion of neuroprogenitor cells, or perturbing adhesive forces as neurons radially migrate, can lead to cortical folding and the formation of gyri and sulci, respectively.

We now provide evidence that balanced retinoic acid (RA) signaling from the meninges is required for maintaining lissencephaly in mice. Loss of *Twist1* via *Sm22a-Cre* in cranial mesenchyme affects cell proliferation in the primitive meninx, resulting in loss of arachnoid fibroblasts that express *Raldh2*, a key enzyme required for RA synthesis (Siegenthaler et al., 2009). *Twist1^FLX/FLX^:Sm22a-Cre* animals develop cortical folds and sulci in the posterior dorsolateral telencephalon, which are rescued by maternal RA supplementation. Notably, this process does not occur via amplification of neuroprogenitor cells, supporting alternative mechanisms for cortical folding and sulci formation as reported in *Flrt1/Flrt3*^−/−^ mutants (Del Toro et al., 2017). Thus, our findings implicate proper meningeal development, downstream from Twist1, and balanced RA signaling for maintaining lissencephaly in mice.

## Results

### *Twist1^FLX/FLX^:Sm22a-Cre* embryos show loss of Twist1 in cranial mesenchyme and arachnoid fibroblasts

To determine the effects of meningeal *Twist1* expression on forebrain development, we conditionally ablated *Twist1* in cranial mesenchyme using *Sm22a-Cre*. *Sm22a-Cre* is active by E10.5 in populations of neural crest and cranial mesoderm that give rise to the three meningeal layers surrounding the forebrain, diencephalon, and cerebellum (El-Bizri et al., 2008; Tischfield et al., 2017). As previously reported, lineage labeling using *R26:Ai14:Sm22a-Cre* revealed *Ai14:td-Tomato* expression in preosteoblasts, dura, and connexin-43(+) arachnoid fibroblasts surrounding the developing forebrain at E14.5, by which time the three meningeal layers are recognized (**Fig. 1A, B**) (Tischfield et al., 2017). By contrast, *Ai14:td-Tomato* expression appeared sparser in developing pia mater underlying the connexin-43+ arachnoid membrane (**Fig. 1C**).

**Figure 1:**
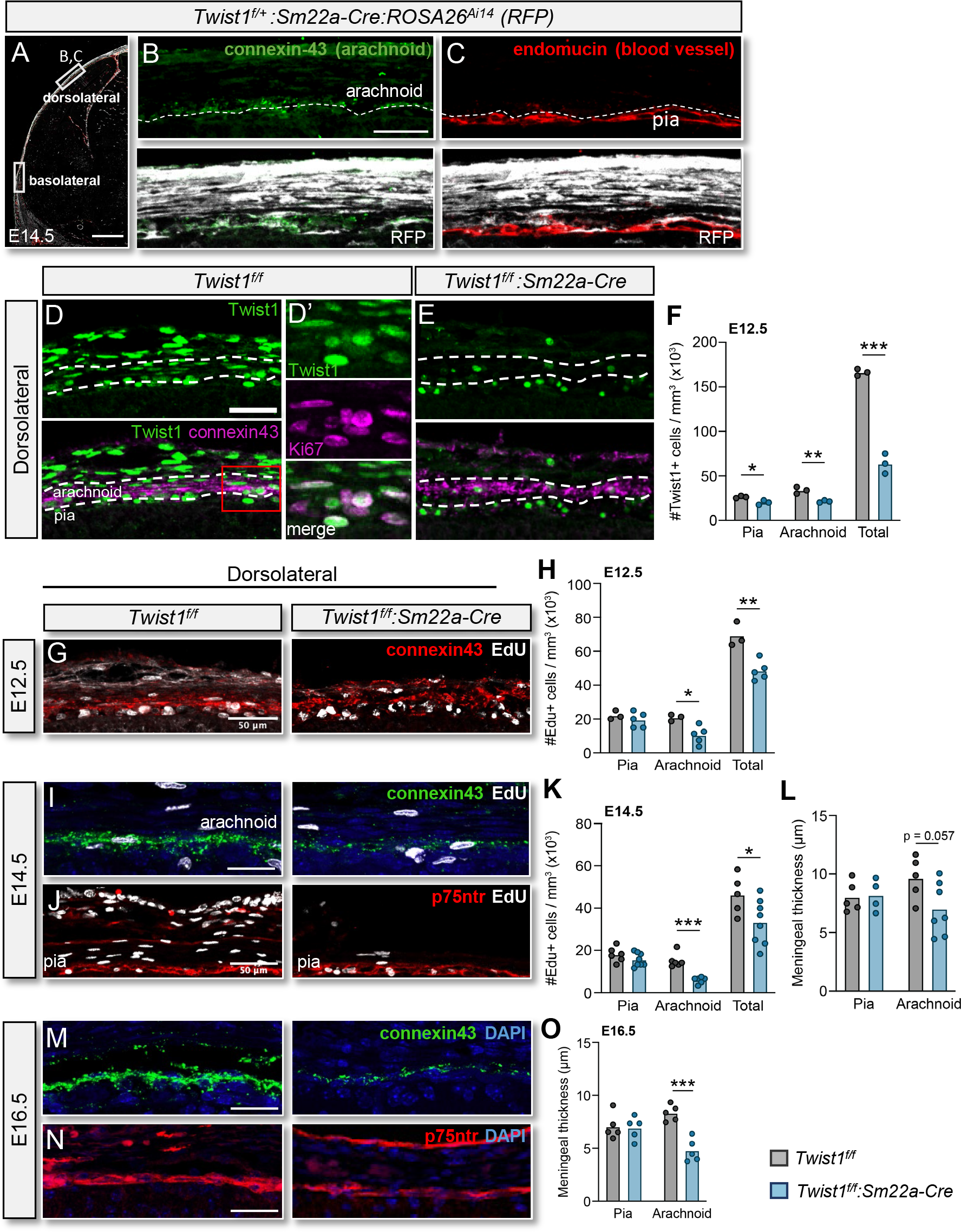
The arachnoid membrane is hypoplastic in *Twist1^FLX/FLX^:Sm22a-Cre* animals. (A-C) *R26:Ai14:Sm22a-Cre* lineage labeling. Cre activity is present in cranial mesenchyme and the arachnoid at E14.5, with sparse activity in the pia. (D) At E12.5, Twist1 is expressed in head mesenchyme and the presumptive arachnoid. Spottier expression is found in pia. The red boxed region (D’) shows Twist1 is expressed in proliferating Ki67+ cells in the arachnoid. (E, F) Twist+ cells are significantly reduced in dorsolateral cranial mesenchyme and presumptive arachnoid in *Twist1^FLX/FLX^:Sm22a-Cre* versus *Twist1^FLX/FLX^* (n=3/genotype). (G, H) At E12.5, *Twist1^FLX/FLX^: Sm22a-Cre* embryos show less EdU^+^ cells in presumptive arachnoid versus *Twist1^FLX/FLX^* (n=3, *Twist1^FLX/FLX^*; n=5 *Twist1^FLX/FLX^:Sm22a-Cre*). (I, K, L) At E14.5, EdU^+^ cells in dorsolateral cranial mesenchyme and arachnoid are reduced and the membrane is hypoplastic (n=5, *Twist1^FLX/FLX^*; n=8, *Twist1^FLX/FLX^:Sm22a-Cre*). (J-L) Pial thickness and the number of EdU+ cells is unaffected in *Twist1^FLX/FLX^:Sm22a-Cre* animals. (M-O) By E16.5, the pial and arachnoid membranes have differentiated, according to p75ntr and connexin-43 labeling, but the arachnoid remains hypoplastic in *Twist1^FLX/FLX^:Sm22a-Cre* embryos. (L, O) Average thickness of the pial and arachnoid membranes (E14.5, n=4/genotype: E16.5, n=6/*Twist1^FLX/FLX^*; n=7/*Twist1^FLX/FLX^: Sm22a-Cre*). One-way ANOVA with Sidak’s multiple comparison test. *\*P<0.05, **P<0.01, ***P<0.001*. Data shows the mean. Scale bars: 500 μm (A), 50 μm (B, D, G, J), 25 μm (I, M, N)

Similarly, Twist1+ cells were found in cranial dermis and the presumptive connexin-43(+) arachnoid membrane at E12.5, but expression was lower in the pia and absent in blood vessels and neurons (**Fig. 1D**). Twist1+ cells also co-localized with proliferating Ki67+ cells in the arachnoid layer (**Fig. 1D’**). In *Twist1^FLX/FLX^:Sm22a-Cre* embryos, the numbers of Twist1+ cells was significantly reduced in cranial mesenchyme surrounding the dorsolateral forebrain, including the differentiating connexin-43(+) arachnoid membrane (**Fig. 1E**). Twist1+ cells were also reduced in cranial mesenchyme surrounding the basolateral forebrain, but to a lesser extent compared to dorsolateral regions (**Fig. S1A-C)**. The number of Twist1+ cells in the pia was also reduced, but not to the extent seen in the arachnoid membrane (**Fig. 1E, F**). Taken together, *Twist1^FLX/FLX^:Sm22a-Cre* produces a model in which Twist1+ cells are lost in the leptomeninges (arachnoid and pia), but more so in the arachnoid membrane and throughout dorsolateral versus basolateral cranial mesenchyme.

### Twist1 regulates cell proliferation in the meninges

We examined cell proliferation in developing cranial mesenchyme and the leptomeninges at E12.5 and E14.5. In E12.5 *Twist1^FLX/FLX^:Sm22a-Cre* mutants, we observed less proliferating cells in regions of interest within the dorsolateral and basolateral cranial mesenchyme following a brief one-hour EdU pulse. Less EdU+ cells were found in the presumptive connexin-43(+) arachnoid layer, whereas EdU labeling was comparable in the pia **(Figs. 1G, H, S1F, G**). By E14.5, Twist1 protein was significantly downregulated in the meninges (**Fig. S1D, E**), but cell proliferation remained decreased throughout dorsolateral and basolateral regions following a one-hour pulse with EdU **(Figs. 1K, S1J)**. The number of EdU+ cells co-labeled with connexin-43(+) was reduced, whereas EdU labeling in the pia, as visualized by p75ntr staining, remained unaffected (**Figs. 1I-K, S1H-J**). The arachnoid membrane was hypoplastic in both basolateral and dorsolateral regions by E14.5, whereas the pial layer, which showed normal amounts of EdU-labeled cells, was unaffected (**Figs. 1L, S1K**). Thus, consistent with *Sm22a-Cre* activity and Twist1 expression, cell proliferation was reduced in cranial mesenchyme giving rise to arachnoid mater, whereas proliferation in the pia was comparable between *Twist1^FLX/FLX^:Sm22a-Cre* mutants and *Twist1^FLX/FLX^* controls. The leptomeninges differentiated by E14.5 and E16.5, as assessed by p75ntr, Connexin-43, and Crabp2 staining, which labels the pia, arachnoid, and both the dura and arachnoid membranes, respectively **(Figs. 1M, N, S1L, M)**(DeSisto et al., 2020). The arachnoid layer remained hypoplastic in E16.5 *Twist1^FLX/FLX^:Sm22a-Cre* mutants (**Figs. 1O, S1N**). These results suggest loss of Twist1 affects cell proliferation and attenuates the expansion of cranial mesenchyme and the meninges, although the meninges continue to differentiate and express layer-specific markers.

### *Twist1^FLX/FLX^:Sm22a-Cre* mutants develop cortical bumps and sulci

Loss of *Twist1* in cranial mesenchyme produced macroscopic cortical abnormalities in postnatal animals. Compared to the smooth cortical surface characteristic of lissencephalic mice, nearly all *Twist1^FLX/FLX^:Sm22a-Cre* mutants showed tapering of the anterior cortex and macroscopic bumps and dips on the cortical surface (**Fig. 2A-C**). A smaller fraction had caudal shortening of the cortex (**Fig. 2B, Table S1**). Tapering of the anterior cortex corresponded to the shape of the skull, which showed coronal synostosis and misshapen frontal bones (**Fig. 2D**). We postulated that macroscopic bumps on the cortical surface (mainly found in the anterior telencephalon) could have resulted from neuronal breaches of the pial basement membrane. However, sections through the anterior cortex of P10 animals revealed cortical layering was intact beneath the bumps as assessed by Satb2 (layers 2-4), Ctip2 (layer 5), and FoxP2 (layer 6) staining **(Fig. 2E-F’’)**. Furthermore, the pial basement membrane was intact at E16.5 and P10 as assessed by Laminin-α1 staining (**Fig. 2G-H’, I**), and nestin+ radial glia endfeet were properly anchored to the pial basement membrane by E16.5 (**Fig. 2G-H”**). Rather, macroscopic bumps on the anterior cortical surface of *Twist1^FLX/FLX^:Sm22a-Cre* mutants appeared to result from focal increases in the numbers of Foxp2 and Ctip2 deep-layer neurons (**Fig. 2E, F**).

**Figure 2:**
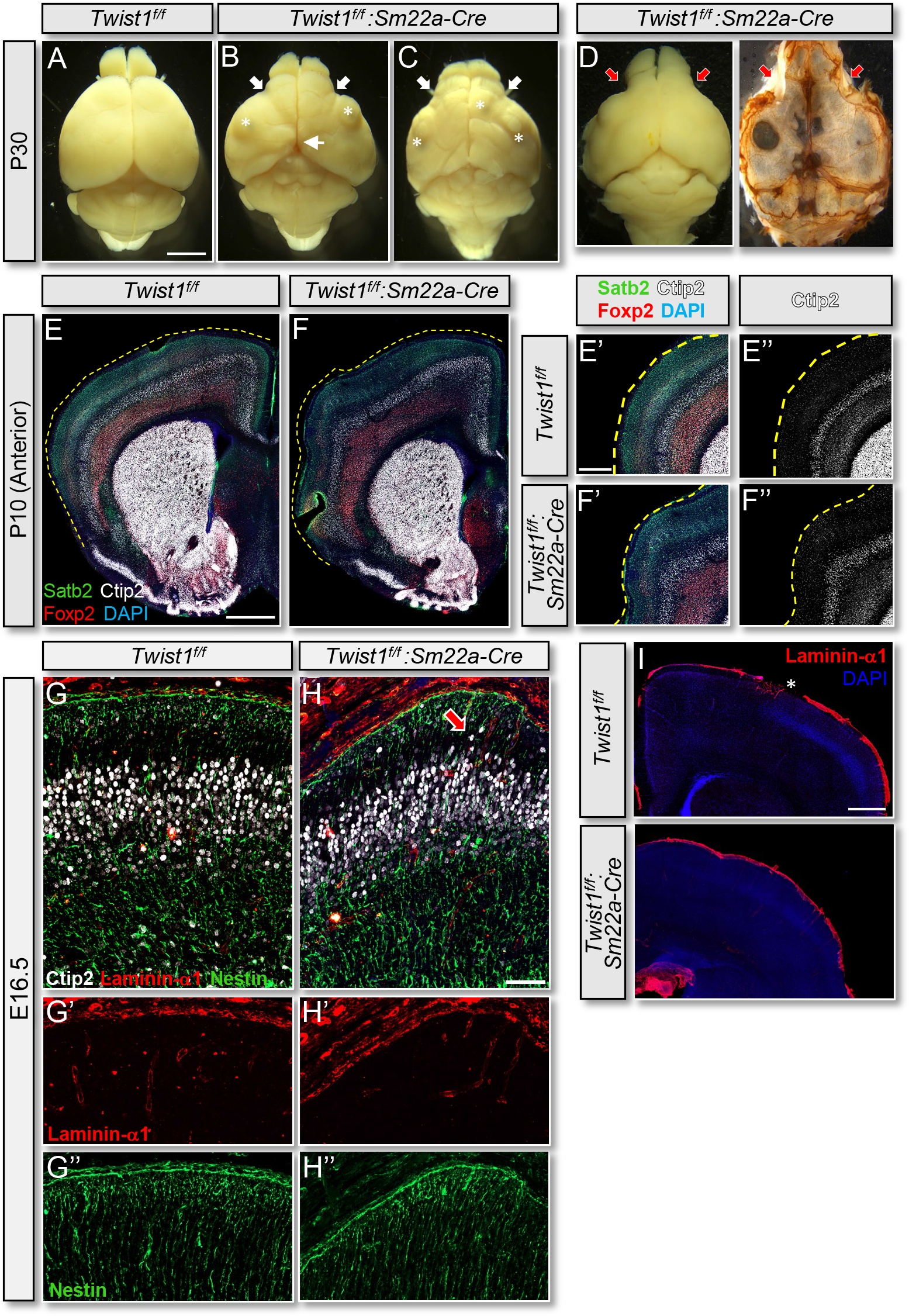
*Twist1^FLX/FLX^:Sm22a-Cre* animals develop macroscopic cortical bumps. (A-C) P30 *Twist1^FLX/FLX^: Sm22a-Cre* brains show narrowing of the anterior cortex (thick arrows), with numerous bumps (asterisks) and dips on the cortical surface. Caudal shortening of the cortex (thin arrow) is present in some. (D) Narrowing of the anterior cortex corresponds to skull abnormalities and narrowing of the frontal bones. (E-F’’) P10 coronal sections through the anterior cortex and corresponding magnified images stained with layer-specific markers Satb2, Ctip2, and Foxp2. *Twist1^FLX/FLX^:Sm22a-Cre* animals show gyri-like bumps with preserved cortical lamination. (G-H”) The pial basement membrane develops normally at E16.5 in *Twist1^FLX/FLX^:Sm22a-Cre* embryos, and nestin+ radial glial endfeet are properly attached. Note that occasional over-migration of Ctip2+ neurons adjacent to developing sulcal regions (arrow in G) occurs alongside normal development of the pial basement membrane and proper attachment of radial glial endfeet. (I) The pial basement membrane overlying cortical bumps is intact at P10. Asterisk indicates sectioning artifact. Scale bar: 500mm (A-C), 1mm (E, E’), 50μm (G, H), 500μm (I)

Strikingly, 44% of *Twist1^FLX/FLX^:Sm22a-Cre* animals showed cortical folding and sulci that persisted into adulthood (**Fig. 3A-D’**). Sulci were typically localized to one cortical hemisphere and were most commonly found in the posterior dorsolateral sensorimotor cortex at or near the hippocampus. Sulci were present with and without cortical invagination (**Fig. 3B’, D’**). Cortical layering was grossly normal and the overlying pial basement was intact (**Fig. 3D-F’**). These telencephalic fissures appeared to be bona fide sulci since they maintain hallmarks of proper cortical folding - folding of the cortical layers and pial surface without disruption of the ventricular surface (Borrell, 2018). Collectively, these findings suggest that loss of Twist1 in cranial mesenchyme and developing arachnoid mater, especially in dorsolateral regions surrounding the posterior telencephalon, produces cortical folding in mice.

**Figure 3:**
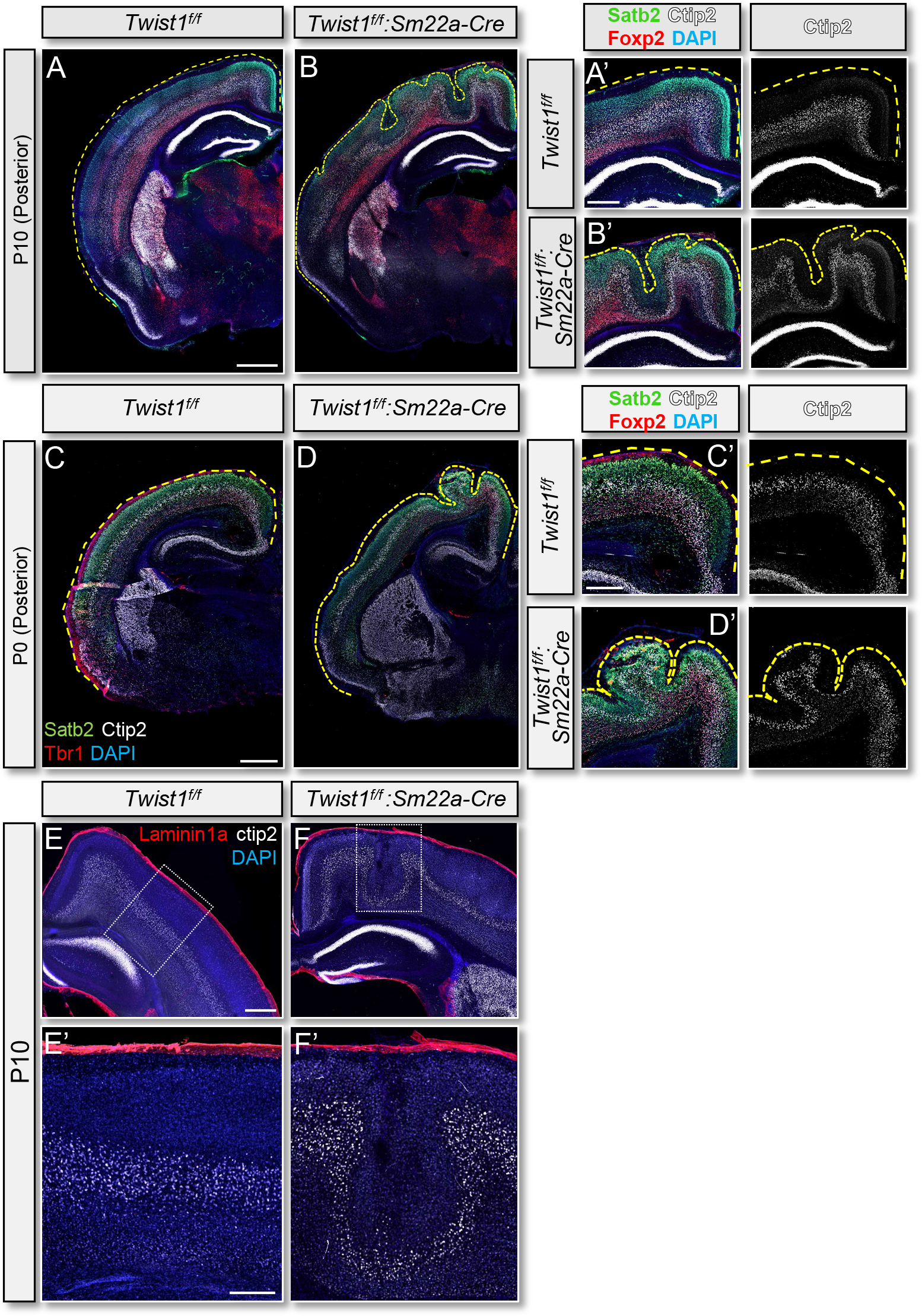
*Twist1^FLX/FLX^:Sm22a-Cre* animals develop cortical folding and sulci. (A-B’) P10 coronal sections through the posterior cortex at the level of the hippocampus. *Twist1^FLX/FLX^:Sm22a-Cre* animals show cortical folding and sulci along the dorsolateral cortex and near the midline. Cortical layers are preserved but collectively fold in these sulci-like structures. The underlying ventricular surface is normal. (C-D’) Sulci are present by P0 in *Twist1^FLX/FLX^:Sm22a-Cre* animals. (E-F’) The pial basement membrane overlying sulci in the posterior cortex is intact at P10. Ctip2 labels layer 5 neurons. Scale bar: 1mm (A, A’), 500μm (C, C’), 250μm (E, E’)

### Cortical folding occurs independent of neuroprogenitor cell amplification

Cortical folding was first apparent at E16.5 in the form of invaginations in the dorsolateral telencephalon that corresponded to the eventual location of sulci (**Fig. 4A-B**). Cortical layering was grossly normal in presumptive sulcal regions without perturbation to the ventricular surface (**Fig. 4A-B**). Although we could detect occasional Ctip2^+^ cells that appeared to migrate further into the cortical plate at regions adjacent to presumptive sulci (**Fig. 2H**), deep layer neurons were not found in superficial layers, or vice versa, as commonly seen in neuronal heterotopias or in situations where radial glia endfeet detach from the pial basement membrane causing radial migration defects (Zarbalis et al., 2007; Myshrall et al., 2012; Inoue et al., 2008). A hallmark of proper cortical folding is varying thicknesses of the cortex and cortical layers in gyral and sulcal regions, with thicker and thinner layers in gyri and sulci, respectively (Borrell, 2018). In agreement, average cortical thickness was reduced and cortical layers were thinner in *Twist1^FLX/FLX^:Sm22a-Cre* sulci compared to adjacent regions and the equivalent region in *Twist1^FLX/FLX^* controls (**Fig. 4C, D**). Cell counts in regions adjacent to sulci in *Twist1^FLX/FLX^:Sm22a-Cre* animals showed comparable numbers of Tbr1+ and Ctip2+ deep layer neurons, but a slight decrease in the number of Satb2^+^ upper layer neurons compared with controls. In sulci, the numbers of Ctip2+ and Tbr1+ deep layer neurons were generally reduced, accompanied by a larger reduction in the number of Satb2^+^ upper layer neurons (**Fig. 4E**). We also found differences in the distribution of projection neurons along the tangential axis, as neurons tended to be located closer to the pial surface by E16.5, especially in sulcal regions (**Fig. 4F-J**).

**Figure 4:**
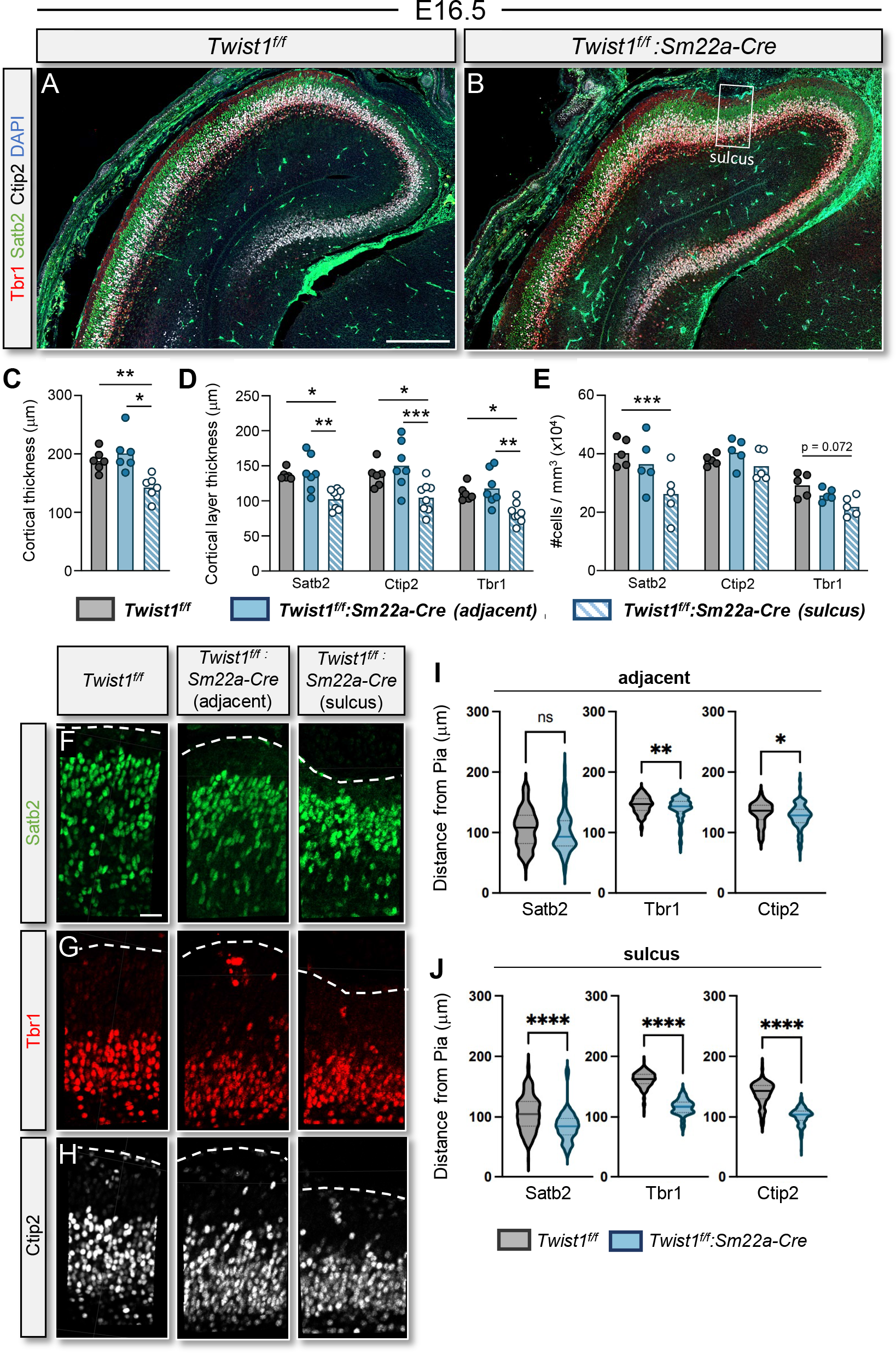
Cortical layering is preserved but the number of neurons in sulcal regions and their relative distance from the pial surface is altered. (A, B) E16.5 coronal sections through the dorsolateral cortex. Tbr1, Satb2, and Ctip2 staining shows cortical layering is grossly normal in *Twist1^FLX/FLX^:Sm22a-Cre* embryos. Sulci are first detected at this timepoint (boxed region). (C) Average cortical thickness is reduced in *Twist1^FLX/FLX^:Sm22a-Cre* sulci (n=6) compared to the adjacent area and *Twist1^FLX/FLX^* animals (n=6). Brown-Forsythe and Welch’s ANOVA test. (D) Average thickness of both the deep (Tbr1, Ctip2) and superficial layers (Satb2) is also reduced in sulci. Two-way ANOVA with multiple comparisons. (E) The numbers of Satb2 cells are significantly reduced in *Twist1^FLX/FLX^:Sm22a-Cre* sulci. One-way ANOVA with multiple comparisons. ( F-H) Representative ROI used to measure distance of Sabt2, Tbr1, and Ctip2 cells relative to the pia. (I, J) Violin plots show that Satb2, Ctip2, and Tbr1 cells located in and adjacent to sulci regions are distributed closer to the pial surface in *Twist1^FLX/FLX^:Sm22a-Cre* cortices. Welch’s T-test. Data shows the mean. *\*P<0.05, **P<0.01, ***P<0.001, ****P<0.0001*. Scale bar: 200μm (A), 30μm (F)

Cortical folding can develop from an accumulation of apical neuroprogenitors (aRGCs) in the VZ, which produce basal neuroprogenitors (bRGCs and bIPs) that give rise to neurons (DelValle-Anton and Borrell, 2022). We investigated whether the number of neuroprogenitor cells differed in sulcal regions at E14.5 and E16.5 by staining for Pax6 (RGCs) and Tbr2 (IPs). We found comparable numbers of Pax6+ RGCs and Tbr2+ IPs in the posterior dorsolateral telencephalon at E14.5, and the numbers of cells co-labeled with EdU were similar (**Fig. 5A-F, I)**. The number of post-mitotic Tbr1+ deep layer neurons was normal and EdU+ cells were not present outside the VZ in the cortical plate (**Fig. 5G-H’, L**) We also found comparable numbers of Pax6+ RGCs and Tbr2+ IPs in the posterior dorsolateral telencephalon at E16.5, in and adjacent to sulcal regions **(Fig 5. J-M**). Additionally, the length of the dorsal forebrain was normal and we still could not detect Pax6+ cells outside the VZ that would be indicative of bRGCs (**Fig. 5N**).

**Figure 5:**
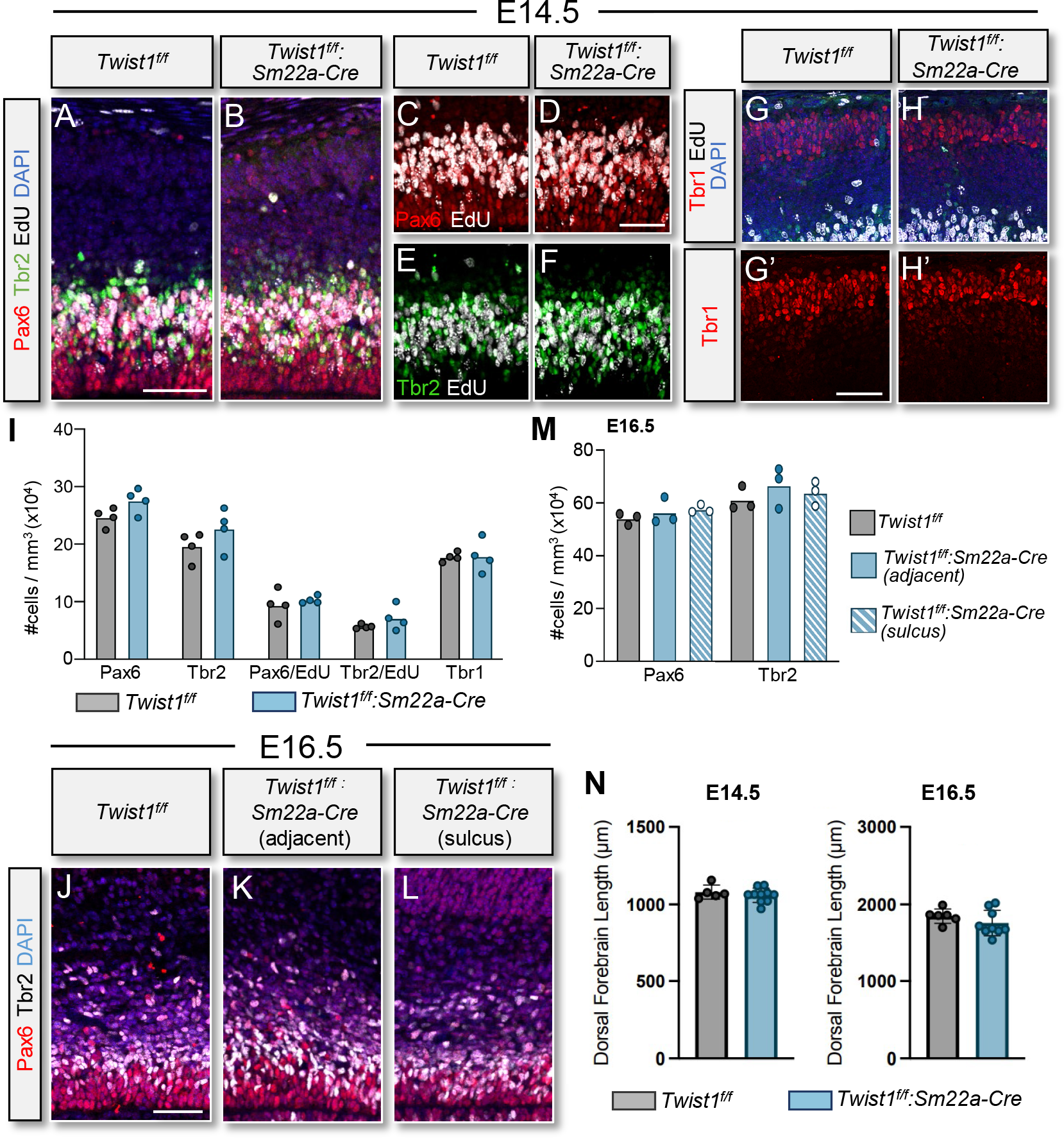
Cortical folding occurs in the absence of neuroprogenitor cell amplification. (A-I) E14.5 dorsolateral cortices and cell counts. In sulcal regions, *Twist1^FLX/FLX^:Sm22a-Cre* embryos show normal numbers of RGCs (Pax6+), IPs (Tbr2+), and cells co-labeled with EdU. Pax6+ and Tbr2+ cells are not found outside the SVZ. (G-I) The numbers of differentiated Tbr1+ neurons at E14.5 are comparable between *Twist1^FLX/FLX^* and *Twist1^FLX/FLX^:Sm22a-Cre* brains, and dividing EdU+ cells are not present in the cortical plate. One-way ANOVA with Sidak’s multiple comparison test. Data shows the mean. N=6 for *Twist1^FLX/FLX^:Sm22a-Cre* and *Twist1^FLX/FLX^* (C, D) and N=5 for both genotypes (E). (J-M) The numbers of Pax6+ RGCs and Tbr2+ IPs are normal at E16.5 in the dorsolateral cortex surrounding sulcal regions, suggesting cortical folding occurs independently of neuroprogenitor cell amplification. One-way ANOVA with Sidak’s multiple comparison test, n=3 for each genotype. (N) The length of the dorsal forebrain is comparable between *Twist1^FLX/FLX^ and Twist1^FLX/FLX^:Sm22a-Cre* embryos at E14.5 and E16.5. N=5 and 10 for *Twist1^FLX/FLX^ and Twist1^FLX/FLX^:Sm22a-Cre*, respectively, at E14.5. N=6 and 9 for *Twist1^FLX/FLX^ and Twist1^FLX/FLX^:Sm22a-Cre*, respectively, at E16.5. One-way ANOVA with Sidak’s multiple comparison test. Scale bar: 50μm (A, D, G’), 30μm (J).

### Raldh2 expression is diminished in the dorsolateral meninges

Cranial mesenchyme and the meninges proper secrete morphogens that regulate cortical development, including retinoic acid (RA) and Bone Morphogenetic Proteins (BMPs) (Siegenthaler et al., 2009; Choe et al., 2012). RA signaling is reported to influence the ability of aRGCs to asymmetrically divide, and later facilitates the transition from multipolar to bipolar morphology required for neurons to radially migrate (Siegenthaler et al., 2009; Haushalter et al., 2017; Choi et al., 2014). Raldh2, an enzyme required for RA synthesis, is produced in arachnoid fibroblasts and is deficient in *Foxc1^hith/hith^* animals, which have hypoplastic meninges and show some phenotypic similarities to *Twist1^FLX/FLX^:Sm22a-Cre* animals. However, *Foxc1^hith/hith^* animals, and especially more severely affected *Foxc1^−/−^* embryos, show pseudo-gyrification with folding of the ventricular surface, breaches in the pial membrane, and cortical layering defects (Siegenthaler et al., 2009). Nonetheless, we asked whether Raldh2 expression and meningeal-derived RA signaling was affected in *Twist1^FLX/FLX^:Sm22a-Cre* embryos. At E14.5, we did not detect a significant decrease in the number of cells expressing Foxc1 in the leptomeninges (**Figs. 6A, B, I, S2A-D, S2L**). We also did not detect loss of Raldh2 expression in the forebrain leptomeninges at this time (**Fig. 6C, D**). Interestingly, at E16.5, the earliest time point when we could detect presumptive sulci, the number of Foxc1+ cells in the dorsolateral - but not basolateral - meninges was significantly reduced (**Figs. 6E, F, J, S2E-L**). Furthermore, we detected focal losses and/or weaker Raldh2 expression in the dorsolateral meninges immediately adjacent to presumptive sulci **(Figs. 6 G-H, S2G-H)**. These results suggest loss of *Twist1* in cranial mesenchyme and the primitive meninx at E12.5 reduces cell proliferation in the arachnoid, leading to eventual loss of Foxc1+ and Raldh2+ cells in the dorsolateral meninges.

**Figure 6:**
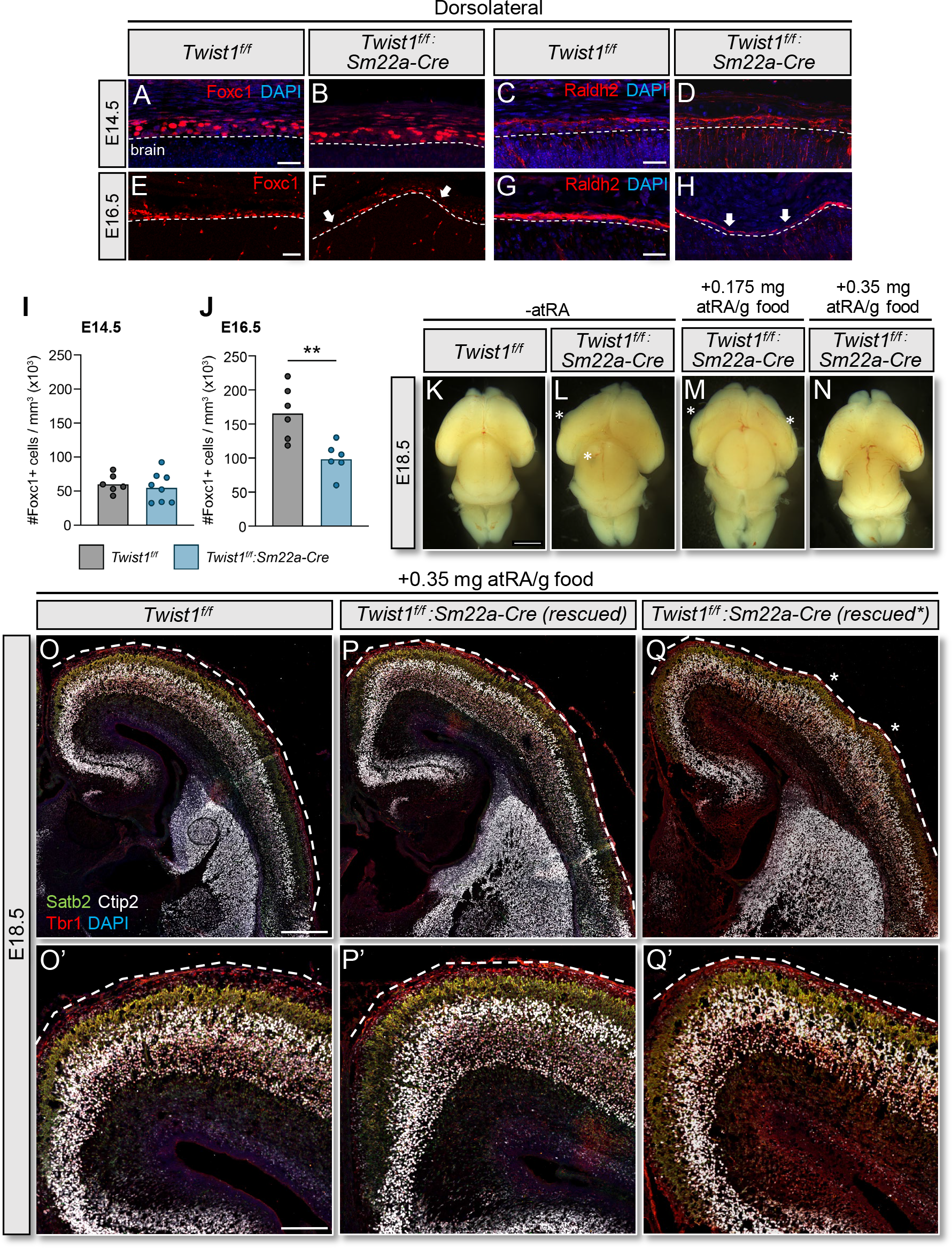
Maternal RA supplementation counteracts loss of leptomeningeal fibroblasts and RA signaling to rescue cortical folding. (A-D, I) At E14.5, the numbers of Foxc1+ leptomeningeal fibroblasts and the levels of Raldh2 staining are normal in *Twist1^FLX/FLX^:Sm22a-Cre* embryos (n=8) compared to *Twist1^FLX/FLX^* (n=6). (E-F, J) At E16.5, the numbers of Foxc1+ leptomeningeal fibroblasts are reduced (arrows) and Raldh2 staining is weaker and discontinuous (arrows) in *Twist1^FLX/FLX^:Sm22a-Cre* embryos (n=6) compared to controls (n=6), particularly surrounding sulcal regions. Welch’s t-test. *\*\*P<0.01*. (K, L) *Twist1^FLX/FLX^* and *Twist1^FLX/FLX^: Sm22a-Cre* E18.5 brains without all-trans retinoic acid (atRA) supplementation. (M, N) *Twist1^FLX/FLX^: Sm22a-Cre* E18.5 brains with atRA supplementation at two doses. (M) Cortical bumps (asterisks) are still visible on *Twist1^FLX/FLX^:Sm22a-Cre* brains at E18.5 with 0.175 mg atRA/g supplementation (n=10). (N) 4/9 *Twist1^FLX/FLX^:Sm22a-Cre* brains supplemented with 0.35 mg atTRA/g food were macroscopically indistinguishable from controls. (O-Q) Coronal sections through the cortex in embryos treated with 0.35 mg atRA/g food. Cortical layering is preserved and no sulci were detected in *Twist1^FLX/FLX^:Sm22a-Cre* brains (n=8). Small cortical bumps (asterisks) are found in a subset of the rescued mutants (5/9), as seen in panel Q (rescued*). (O’-Q’) Magnified regions where sulci are normally found in P0 *Twist1^FLX/FLX^:Sm22a-Cre* brains. Scale bar: 25μm (E, G), 2mm (K), 400μm (O), 200μm (O’).

### Maternal supplementation with RA partially restores lissencephaly in *Twist1^FLX/FLX^:Sm22a-Cre* animals

Localized loss of Raldh2 expression in the dorsolateral meninges prompted us to ask whether maternal supplementation of RA during pregnancy could restore lissencephaly in *Twist1^FLX/FLX^:Sm22a-Cre* animals. We supplemented pregnant dams with either 0.175mg or 0.35mg of RA/gram of food from E12.5 to E17.5 before collecting embryos at E18.5. Supplementing 0.175mg of RA/gram of food resulted in macroscopic bumps and dips on the cortical surface in all mutant embryos (n=10), similar to untreated *Twist1^FLX/FLX^:Sm22a-Cre* animals **(Fig. 6K-M)**. However, supplementing with 0.35mg of RA/gram of food resulted in 44% of embryos (4/9) that, macroscopically, were indistinguishable from controls **(Fig. 6N)**. The remaining five embryos still displayed a bump on the anterior telencephalon **(Fig. 6M)**. The cortex was smooth and shaped normally in rescued embryos and also did not show signs of caudal shortening as observed in some *Twist1^FLX/FLX^:Sm22a-Cre* animals. Importantly, sections through the posterior telencephalon did not reveal the presence of sulci in any of the mutant embryos supplemented with 0.35mg of RA/gram of food **(Fig. 6O-Q’)**. These results suggest regionalized loss of RA signaling during late stages of cortical neurogenesis (i.e., E16.5) is at least partly responsible for the development of sulci in *Twist1^FLX/FLX^:Sm22a-Cre* animals.

### Reducing BMP4 expression in cranial mesenchyme does not increase cortical folding

Downregulation of BMP4 expression in the cranial mesenchyme and neuroepithelium at the dorsal midline of *Lmx1a^−/−^:Lmx1b^−/−^* mutant embryos causes cortical folding by E15.5. This results from increased neurogenesis due to altered Wnt/β-catenin signaling and a significant expansion of Pax6+ bRGCs and Tbr2+ IPs (Chizhikov et al., 2019). Although we do not see expansion of Pax6+ bRGCs or Tbr2+ IPs at E14.5 and E16.5 in *Twist1^FLX/FLX^:Sm22a-Cre* embryos, we previously reported reduced BMP4 expression in the meninges (Tischfield et al., 2017). Thus, we asked if further reducing the levels of BMP4 expression in the meninges, and also in dorsal midline neuroepithelium, would increase the prevalence of cortical folding in the dorsolateral telencephalon. We intercrossed *Twist1^FLX/FLX^: Sm22a-Cre* mutants with a *Bmp4^CFP^* knock-in reporter allele that is haploinsufficient for *Bmp4* (Jang et al., 2010). Reducing the levels of *Bmp4* expression led to the formation of shallow sulci in the anterior telencephalon in some animals and produced more dysmorphic brains, but we did not see an overall increase in the prevalence or extent of cortical folding in the posterior telencephalon. This suggests attenuation of BMP4 signaling is not a primary driver of the phenotype, and loss of RA signaling exerts stronger effects (**Fig. S2M-O’**).

## Discussion

### Evolution of cortical folding in mammals

Cortical folding evolved in large part from increased levels of neurogenesis in the mammalian neocortex. Symmetrical division of aRGCs is prolonged in gyrencephalic mammals, generating more apical neuroprogenitors and, eventually, more bRGCs and bIPs that give rise to neurons (Borrell, 2018; Chizhikov et al., 2019). Interestingly, the amount of bRGCs in marmosets, which have near lissencephalic brains, is similar compared to human and ferret. This suggests that an overall abundance of bRGCs is necessary, but not sufficient, for cortical folding and other factors such as cell cycle duration and the capacity of bRGCs and/or bIPs to self-renew also dictate folding (Kelava et al., 2012). It was previously assumed that larger gyrencephalic animals evolved from smaller lissencephalic ancestors (Welker, 1990). However, evidence suggests that some mammals actually underwent the reverse transition to reacquire lissencephaly (i.e., secondary lissencephaly), as is the case with marmosets and mice who had gyrencephalic ancestors (Kelava et al., 2012; Kelava et al., 2013). Also, bRGCs are rare and bIPs have limited ability to self-renew in mice (Wang et al., 2011; Shitamukai, Konno, and Matsuzaki, 2011). Thus, increasing the overall abundance of neuroprogenitors and their capacity to self-renew and generate neurons is integral to cortical folding, whereas the reverse is true for promoting secondary lissencephaly.

### Role of meningeal Twist1 expression for maintaining secondary lissencephaly in mice

Conditional inactivation of *Twist1* in cranial mesenchyme and the primitive meninx via *Sm22aCre* induces the formation of sulci, which are most often unilateral and located in the posterior dorsolateral telencephalon. Sulci display hallmarks of bona fide cortical folding with preserved cortical lamination and variation in layer thickness between sulci and adjacent regions, without disrupting the underlying ventricular surface. Interestingly, sulci in *Twist1^FLX/FLX^:Sm22a-Cre* animals correspond to conserved sulcal regions in gyrencephalic animals; 40% showed folding near the corresponding intraparietal sulcus and 13% in the Sylvian sulcus. One animal had folding in both locations. The closest gyrencephalic small rodent relative, the guinea pig, has primitive folding in the intraparietal and Sylvian sulci (Hatakeyama, Sato, and Shimamura, 2017). Overall, cortical folding patterns are generally conserved in mammals, although they can differ across species (Welker, 1990). Repeated occurrences of folding in areas corresponding to the intraparietal and Sylvian sulci in transgenic mice and guinea pigs suggests that some cortical regions are more prone to folding.

Our findings show some similarities, but notable differences, to mouse transgenic models with induced cortical folding and gyri formation. Knockdown of *Trnp1* causes gyri formation and folding in the intraparietal sulcus (Stahl et al., 2013), and overexpression of FGF2 causes folding near the Sylvian sulcus in the rostrolateral cortex during embryogenesis (Rash et al., 2013). Similar to *Lmx1a^−/−^:Lmx1b^−/−^* mutants (Chizhikov et al., 2019), cortical folding and gyri formation in these models is thought to occur via amplification of neuroprogenitor cells. By contrast, these findings are absent from sulcal regions in *Twist1^FLX/FLX^:Sm22a-Cre* animals. We note, however, that further studies are necessary to determine the etiology of macroscopic bumps in the anterior telencephalon, which are morphologically distinct from sulci and appear to involve focal increases in the numbers of deep layer neurons.

*Flrt1^−/−^/3*^−/−^ mice develop sulci in both the Sylvian and intraparietal sulcus, and show similar spatiotemporal development of sulci compared with *Twist1^FLX/FLX^:Sm22a-Cre* animals (Del Toro et al., 2017). Notably, cortical folding in *Flrt1^−/−^/3^−/−^* mice is also independent from neuroprogenitor amplification. Instead, these animals show changes to the clustering and migration speed of neurons, leading to progressive cortical folding and the formation of sulci (Del Toro et al., 2017). These findings agree with differential tangential expansion theory; regions with high neuronal density expand less compared to regions with lower density, and mechanical instability resulting from differential expansion leads to “buckling” of the cortex (Ronan et al., 2014; Garcia, Kroenke, and Bayly, 2018). Thus, in *Flrt1^−/−^/3*^−/−^ mice, abnormal neuronal clustering promotes mechanical instability and cortical folding by forming areas of high density (clusters of fast migrating neurons) and low density (neurons with normal tangential distribution and migrating speed). Interestingly, we find that projection neurons at sulcal regions are distributed closer to the pial surface at E16.5, suggesting aspects of their radial migration, such as speed, may be altered. Future experiments are needed to address whether neuronal migration speed is differentially affected in sulcal and adjacent regions in *Twist1^FLX/FLX^:Sm22a-Cre* animals, and whether this affects neuronal density leading to sulci.

Our findings suggest that *Twist1* maintains secondary lissencephaly in mice, at least in part, through expansion of arachnoid fibroblasts in the dorsolateral meninges which produce RA. In *Twist1^FLX/FLX^:Sm22a-Cre* animals, we find less proliferating arachnoid fibroblasts (i.e., Foxc1 cells) and focal loss of Raldh2+ cells at E16.5 overlying presumptive sulci. Considering cortical folding is not reported in conditional *Raldh2* null mutants (Haushalter et al., 2017), how might loss of RA signaling contribute to sulci formation in *Twist1^FLX/FLX^:Sm22a-Cre* animals? Conditional loss of *Raldh2* expression between E12.5-E17.5 affects radial neuron migration, causing some neurons to stall because the transition from multipolar to bipolar morphology is affected (Haushalter et al., 2017; Choi, Park, and Sockanathan, 2014). Thus, we speculate that regionalized loss of Raldh2 may differentially impact radial migration, such that populations deficient in RA signaling may migrate slower than adjacent regions receiving adequate RA. Cortical folding would therefore be absent in conditional *Raldh2* null models, in which RA signaling is uniformly downregulated across the cortex, rather than localized loss to regions overlying sulci in *Twist1^FLX/FLX^:Sm22a-Cre* animals, which may alter neuronal clustering. In conclusion, our findings now suggest the meninges can signal to suppress cortical folding and maintain lissencephaly via balanced RA signaling.

---

**Table 1: Cortical phenotypes present in *Twist1^FLX/FLX^:Sm22a-Cre* animals**. Total fraction for each cortical abnormality at P0, P10, and/or P30. Excluding P0, all *Twist1^FLX/FLX^:Sm22a-Cre* brains show narrowing of the anterior cortex (25/25). *Twist1^FLX/FLX^:Sm22a-Cre* brains at all timepoints have ectopic bumps on the dorsal cortical surface, more pronounced at later stages (31/31). Almost half of *Twist1^FLX/FLX^:Sm22a-Cre* brains show cortical folding near the midline (14/31). Some P30 brains have caudal shortening of the cortex (4/10).

**Figure S1: Basolateral forebrain cranial mesenchyme and leptomeninges are more mildly affected in *Twist1^FLX/FLX^:Sm22a-Cre* embryos**. (A-C) At E12.5, the numbers of Twist1+ cells are reduced in basolateral cranial mesenchyme, but are unaffected in the leptomeninges. (D, E) Twist1 expression progressively declines in cranial mesenchyme and is mainly localized to dermis at E14.5 and E16.5, with minimal expression in the Crabp2+ leptomeninges. (F, G) At E12.5, *Twist1^FLX/FLX^:Sm22a-Cre* embryos show reduced numbers of EdU^+^ proliferating cells in basolateral cranial mesenchyme and the presumptive arachnoid membrane, respectively. (H-K) Around the basolateral forebrain at E14.5, the numbers of EdU^+^ proliferating cells are normal in the pia (labeled by p75ntr) but reduced in cranial mesenchyme and the connexin43+ arachnoid membrane, which is hypoplastic. (L-N) At E16.5, the basolateral leptopmeninges have differentiated in *Twist1^FLX/FLX^:Sm22a-Cre* embryos as judged by proper localization of connexin-43 and p75ntr. Average thickness of the pial membrane is unaffected, but the arachnoid membrane is hypoplastic. One-way ANOVA (Sidak’s multiple comparison test). *\*P<0.05, **P<0.01, ***P<0.001*. Data shows the mean, and whiskers show the minimum and maximum data points. Scale bars: 50 μm (A, D), 25 μm (C, F-O).

**Figure S2: Knocking-down Bmp4 expression in *Twist1^FLX/FLX^:Sm22a-Cre* newborns with meningeal RA deficiency does not increase the incidence of cortical folding**. (A-B) Overview images of Foxc1 expression at E14.5, showing comparable expression between *Twist1^FLX/FLX^* and *Twist1^FLX/FLX^:Sm22a-Cre* embryos. (C-F) Foxc1 expression at E14.5 and E16.5 in the basolateral forebrain leptomeninges is normal in *Twist1^FLX/FLX^: Sm22a-Cre* embryos. (G, H) Overview images showing Foxc1 expression at E16.5. Arrows point to regional loss of Foxc1 expression in the dorsolateral meninges. (I, J) Overview images of Raldh2 expression at E16.5. Raldh2 expression appears downregulated and/or discontinuous in the dorsolateral meninges (arrows). (K, L) Quantifications showing the numbers of Foxc1+ cells in the basolateral region at E14.5 and E16.5 (E14.5, n=4 and 6 for *Twist1^FLX/FLX^ and Twist1^FLX/FLX^:Sm22a-Cre*, respectively; E16.5 (n=6/genotype). Welch’s t-test. (M-O’) P0 newborn *Twist1^FLX/FLX^:Sm22a-Cre:Bmp4^CFP/WT^* brains show a similar incidence of sulcal formation as *Twist1^FLX/FLX^:Sm22a-Cre* animals (4/9; 44% penetrance). (M-M’) Coronal sections through the anterior cortex of *Twist1^FLX/FLX^: Sm22aCre:Bmp4^CFP/WT^* newborns shows an increase in the numbers of small dips along the cortical surface (arrows), with corresponding waviness of the Ctip2 layer. (N-N’) More posterior coronal sections through the brains of *Twist1^FLX/FLX^:Sm22a-Cre:Bmp4^CFP/WT^* newborns show similar incidences of cortical folding near the midline compared to *Twist1^FLX/FLX^:Sm22a-Cre* brains. (OO’) In addition to sulci formation near the midline and in the cingulate cortex, sections through the posterior cortex also show increased laminar disorganization and waviness (thin arrows). Scale bar: 200 μm (A, H, J), 25 μm (F), 500 μm (M-O).

## Materials and Methods

### Animals

The following transgenic mouse lines were used: *Twist1*^FLX^ (RRID: MMRRC_016842-UNC), Rosa26:Ai14^tdTomato^ (RRID:IMSR_JAX:007914), Bmp4^CFP^ (Jang et. al, 2010), Sm22a-Cre (RRID:IMSR_JAX:017491). Male and female mice were included for all experiments. Mice were maintained on a mixed background (C57Bl/6: FVB). Embryos obtained from timed matings were considered 0.5 days old upon observance of a plug. The ages of animals in this study include embryonic day (E) 12.5, 14.5 16.5, 18.5 and postnatal days (P) 0, 10 and 30. Experiments were approved and carried out under IACUC protocol PROTO201702623 (M.A.T.).

### Immunohistochemistry

For postnatal timepoints, P10 and P30 mice were anesthetized with ketamine/xylazine and perfused with PBS followed by 4% paraformaldehyde (PFA). For P0 timepoint, mice were decapitated and brains were dissected out and post-fixed in 4% PFA overnight at 4°C. Tissue was embedded in 3% agarose and coronal sections at 60 μm thickness were collected using Leica VT1000S vibratome. For embryonic samples, pregnant dams were checked for plugs daily. For EdU injections, the pregnant dam was injected with 0.002 of EdU, mixed with 1 mL of saline at E12.5 or E14.5. One-hour post-injection, the pregnant dams were sacrificed and embryos were collected and decapitated. Embryonic heads were incubated with PBS with 0.5% heparin, with continuous shaking for 10 minutes at room temperature (RT) before overnight fixation in 4% PFA at 4°C. Fixed tissue was cryoprotected in 30% sucrose, and 20 μm sections were collected using a Leica CM1950 cryostat. Matched sections were antigen retrieved in boiled citrate buffer (pH 6.0) prior to staining.

### Antibodies

Antibodies were diluted in PBST with 0.1% triton and 5% normal goat serum and applied overnight at RT. The following antibodies were used: rabbit anti-Pax6 (1:100, Novus), rabbit anti-Tbr1 (1:100, Abcam), rat anti-Ctip2 (1:1000, Abcam), mouse anti-Satb2 (1:50, Abcam), rabbit anti-*Foxc1* (1:50, Cell Signaling), mouse anti-*Twist1* (1:50, Santa Cruz), mouse antiCrapb2 (1:200, Millipore), rabbit anti-p75ntr (1:200), chicken anti-nestin (Novus, 1:500), rabbit anti-Laminin1a (1:1000), rat anti-Tbr2/EOMES (1:300, Invitrogen), rabbit anti-Raldh2 (1:500, Sigma Aldrich), mouse anti-connexin 43 (1:200, Santa Cruz), rabbit anti-connexin 43 (1:1000, Abcam), rat anti-Ki67 (1:100, Thermo Fisher), rat anti-Endomucin (1:200, Abcam), and rabbit anti-RFP (1:1000, Rockland). For secondaries, goat anti-rat Alexa Fluor 647, goat anti-rabbit Alexa Fluor 546, and goat anti-mouse Alexa Fluor 488 (all 1:1000, Thermo Fisher) were used. EdU labeling was performed according to manufacturer’s instructions using the Click-iT Plus EdU Alexa Fluor 488 kit.

### Microscopy

Image data were acquired using a 10x 0.45NA objective for E18.5, P10, and P30 samples and 20x 0.80NA and 40x (1.4NA) objectives for E12.5, E14.5, and E16.5 samples on a Zeiss LSM800 confocal microscope, with a z-step size of 5 μm. Images were analyzed using Zen software and ImageJ. Cell counts were performed in a sampling window of 200 μm x 100 μm for E12.5 samples and 300 μm x 100 μm for E14.5 and E16.5. Graphing and statistical testing were done in Graphpad Prism 9.

### Analysis

For E12.5 samples, two consecutive sections of 20 μm thickness were used for quantification. EdU or *Twist1* cells were quantified within a 200 μm by 100 μm window in the dorsolateral or basolateral region. To differentiate cells in the leptomeninges, cells that overlapped with connexin-43 staining were considered arachnoidal cells, and those directly below the connexin43 signal were considered pial cells. Sidak multiple comparison tests were performed in GraphPad Prism. For E14.5 samples, two consecutive sections of 20 μm thickness were used for quantification. EdU or *Foxc1* cells were quantified within a 300 μm by 100 μm window in the dorsolateral or basolateral region. EdU^+^ cells were considered pial or arachnoidal cells if they co-localized with p75ntr or connexin-43, respectively. Cells above the arachnoid staining but below the top p75ntr band were counted towards “total meningeal cells”. For Pax6/Tbr2/Tbr1 quantification, three consecutive sections of 20 μm thickness were used for quantification, with a counting window of 250 μm by 100 μm in the dorsolateral telencephalon. For E16.5 samples, three consecutive sections of 10 μm thickness in the dorsolateral area were used for quantification. Gyrus and sulcus counts depended on the location of the gyrus and sulcus, with the counting window being set to 300 μm by 100 μm. Only cells in the cortical plate were counted. Nearest neighbor analysis was done in the same sampling window. For distance measurements from the pia, a pial midpoint in the ROI window was marked as a reference and all measurements were calculated from this point. For cortical thickness measurements, three measurements per ROI and layer were taken and averaged per sample, and the measurements spanned from the bottom of the cortical plate to the top most cell. in the dorsolateral lateral area were used to include quantification for the sulcus and an adjacent area in the mutant sample, and a corresponding area in the control. The counting window is 250 μm by 100 μm. For the cortical length, three consecutive sections were measured starting at the dorsal midline, along the ventricular zone for E14.5 and along the bottom of the cortical plate for E16.5, and the measurements were averaged per sample. For meningeal thickness measurements in the dorsolateral and basolateral regions, the shortest distance that is orthogonal to the pia (labeled by p75ntr) and the arachnoid (labeled by connexin-43) was measured three times, and the averages recorded.

### Retinoic Acid Rescue

Pregnant dams were removed from the mating cage 10 days past observance of the plug and then weighed daily. Upon separation, food pellets were removed and replaced with 0.3 g of cherry flavored Nutra-Gel diet packs (Bio-Serv) in an opaque plastic container. Food was replenished daily. At E12, the Nutra-Gel was supplemented with all-trans retinoic acid (atRA, Sigma R2625-100MG). For 0.175 mg atRA/g food, 0.002g of atRA was dissolved with 1 mL of corn oil and mixed with 12g of Nutra-Gel. For 0.35 mg atRA/g food, 0.004g of atRA was dissolved with 2 mL of corn oil and mixed with 12 g of Nutra-Gel. All preparations involving atRA were done in the dark, and the mice were fed daily at 4pm. Pregnant females consumed ~0.3 g of Nutra-Gel with atRA daily until the morning-noon at day 18, at which point the female dam was euthanized and the pups were collected. Whole brain tissue was dissected and incubated in 4% PFA overnight at 4°C before being washed with PBS and sunk in 10% sucrose for cryo-embedding.

## Acknowledgments

The authors thank Kush Desai for technical assistance and also Marianne Polunas who provided assistance from the Rutgers Research Pathology Services Core.

## Author contributions

Conceptualization: M.A.T; Formal analysis: K.T.H, S.S; Data curation: K.T.H, P.S.A, M.J, M.J.M; Writing-original draft: M.A.T; Writing-review & editing: M.A.T, K.T.H., M.J.M; Supervision: M.A.T.; Funding acquisition: M.A.T.

