## Supplementary figures and images for "Twist1 and balanced retinoic acid signaling act to suppress cortical folding in mice"

### Supplemental Figure 1

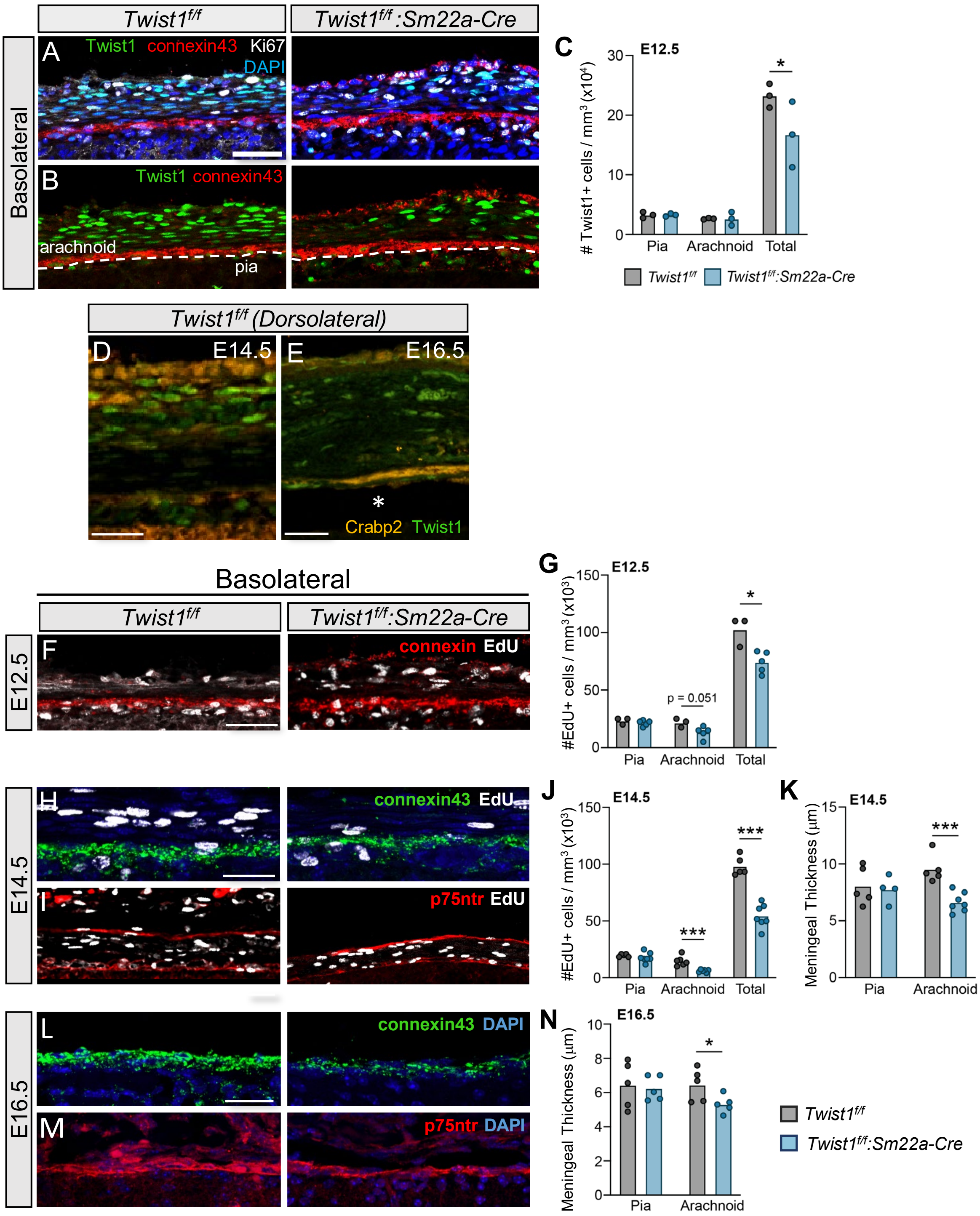

### Supplemental Figure 2

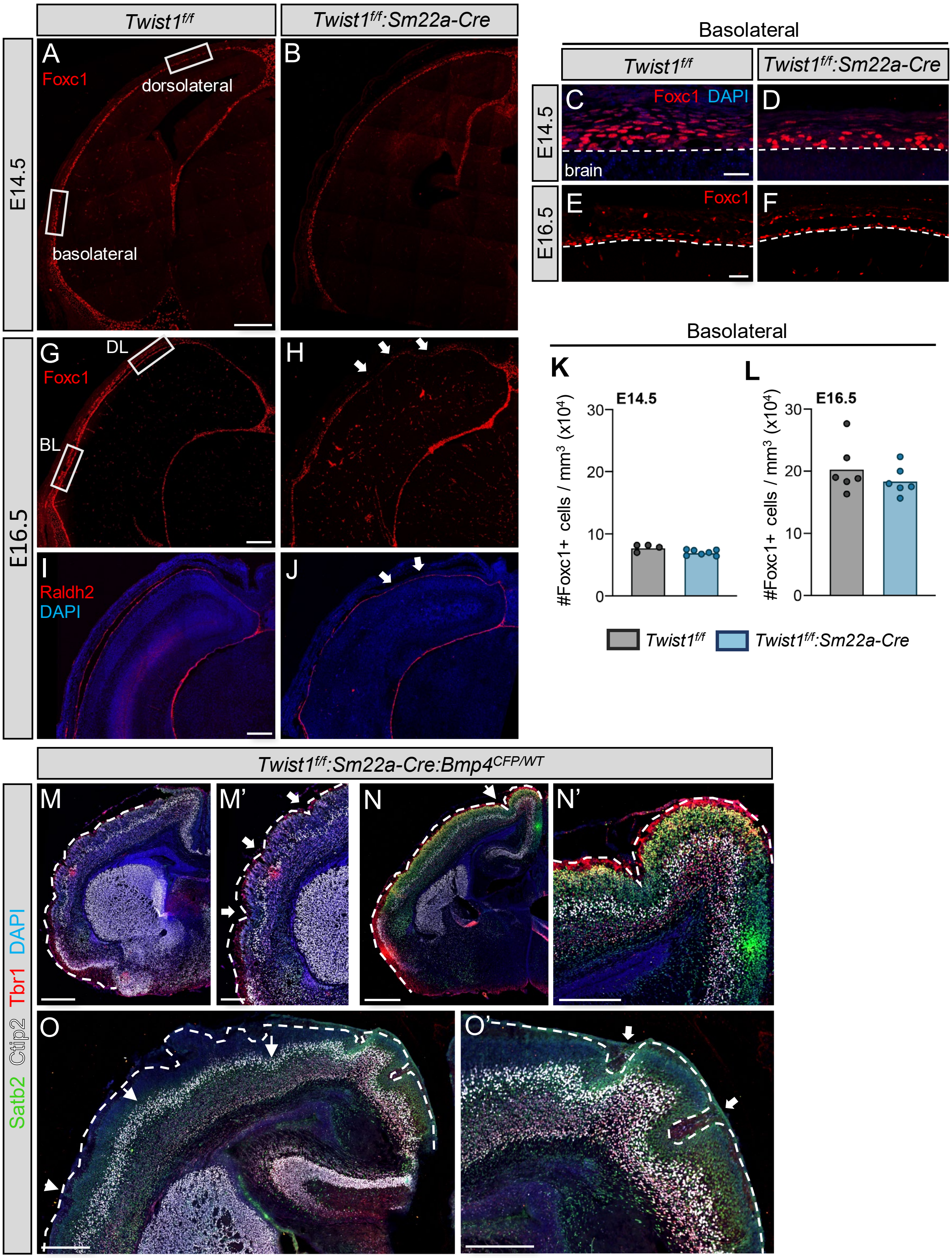

### Supplemental Table 1

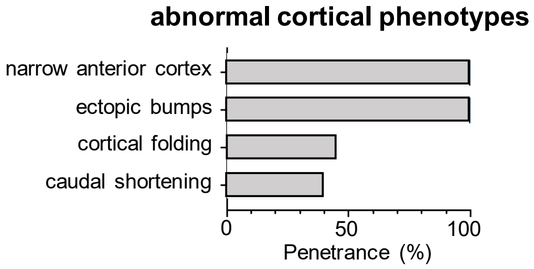
